# Neuro-mesodermal assembloids recapitulate aspects of peripheral nervous system development *in vitro*

**DOI:** 10.1101/2022.06.30.498240

**Authors:** Anna F. Rockel, Nicole Wagner, Süleyman Ergün, Philipp Wörsdörfer

## Abstract

Here we describe a novel neuro-mesodermal assembloid model which recapitulates aspects of peripheral nervous system (PNS) development such as neural crest cell (NCC) induction, delamination, migration and sensory as well as sympathetic ganglion formation. The ganglia send neuronal projections to the mesodermal as well as the neural compartment. Axons in the mesodermal part are associated with Schwann cells. In addition, peripheral ganglia as well as nerve fibers interact with the co-developing vascular plexus, forming a neurovascular niche. Finally, developing sensory ganglia show response to capsaicin treatment indicating their functionality.

The presented assembloid model could help to uncover mechanisms of NCC delamination, migration and PNS development in the human tissue context. Moreover, the model could be used for toxicity screenings or drug testing. The co-development of mesodermal and neuroectodermal tissues and of a well-organized vascular plexus along with a peripheral nervous system allows to investigate the crosstalk between neuroectoderm and mesoderm and between peripheral neurons/neuroblasts and endothelial cells. Such interactions influence NCC delamination and migration, sensory neuron differentiation and rearrangement of the primitive vascular plexus in the embryo.

## Introduction

Within the last decade, numerous human stem cell-based 3D cell culture models were explored, mimicking the development of different organs and tissues. These tissue models are termed organoids. Organoids can provide an unprecedented opportunity to investigate and understand fundamental aspects of human developmental biology *in vitro* ^1^. To increase tissue complexity and allow multi-lineage interaction, different organoids were combined forming assembloids ^2^. In addition, so-called embryoids were established ^3^. Although, most organs and tissues have been modeled as organoids or assembloids, suitable 3D *in vitro* models of peripheral nervous system development are still rare.

The nervous system is divided into two parts, the central and the peripheral nervous system (PNS) which closely interact with each other. The PNS develops mostly from a highly fascinating cell population, the so-called neural crest cells (NCCs) ^4^. NCCs are of ectodermal origin and arise at the neural plate border. They undergo an epithelial to mesenchymal transition (EMT), start to delaminate and finally migrate into distinct regions of the embryo shortly after neurulation. During this process, a part of the NCC population undergoes natural reprogramming processes involving Oct4 reactivation to broaden their differentiation potential ^5^. As soon as migrating NCCs reach their destination, cells start to differentiate e.g., into sensory or sympathetic neurons, Schwann cells, melanocytes as well as smooth muscle cells, chondrocytes, or osteocytes.

Here we describe a neuro-mesenchymal assembloid model which recapitulates peripheral nervous system development *in vitro*. The assembloid model allows neuro-mesodermal interactions and is suitable for the investigation of neural crest cell induction, delamination, migration, and early peripheral nervous system development.

## Results

To generate the assembloid model, iPS cell derived Sox1^+^ neuroepithelial cell aggregates (NT) and mesodermal cell aggregates (MT) were brought in co-culture (Fig. 1 A-B). The purpose of this co-culture is to generate a neuro-mesodermal interface, at which an intercellular crosstalk between both tissue compartments takes place. Neuro-mesodermal interactions were discussed to trigger neural crest cell induction and delamination in the embryo. Moreover, the mesenchymal part provides guidance for migrating NCCs and growing peripheral axons ^6^.

**Figure 1:**
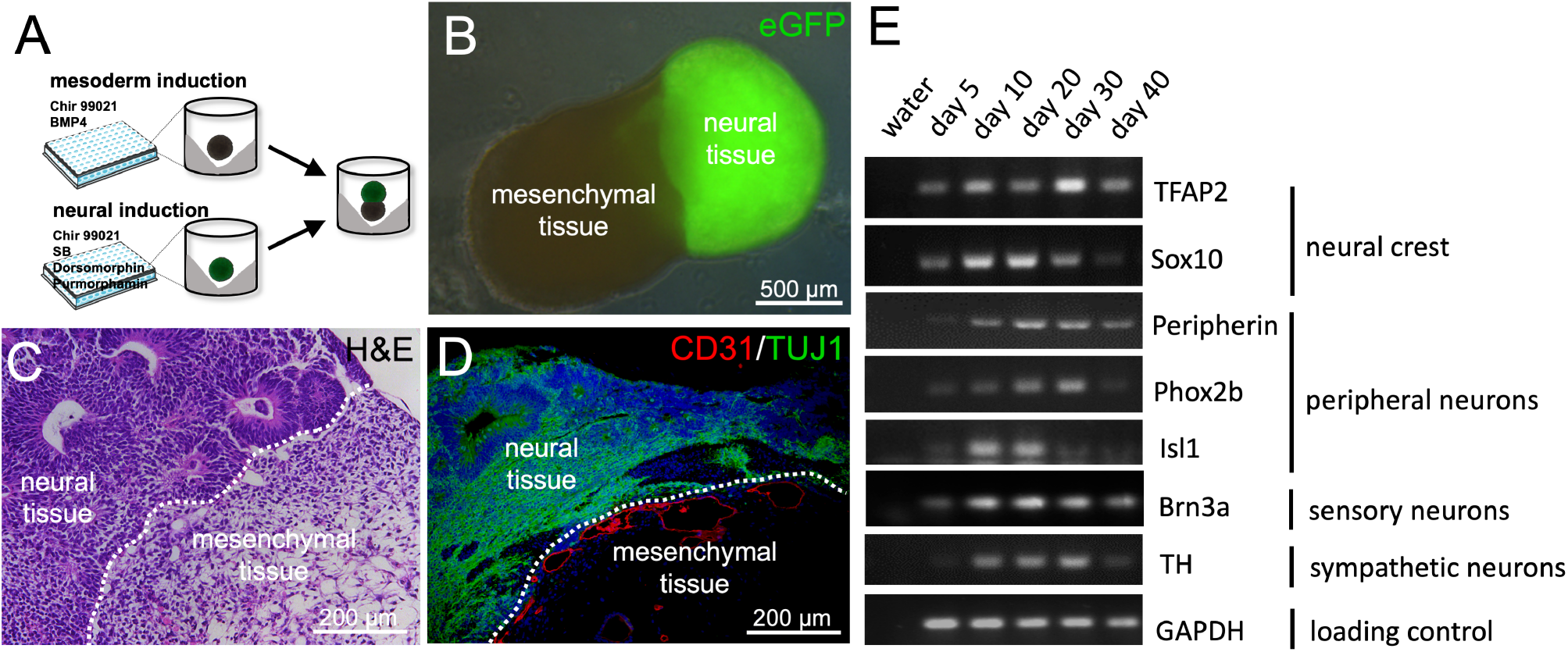
Generation and initial characterization of neuro-mesodermal assembloids. **A:** Schematic representation of the experimental workflow. **B:** Depiction of a typical neuro-mesodermal assembloid at day 14 of co-culture. The neural organoid was generated using GFP-labeled iPS cells. **C:** Histological section of a neuro-mesodermal assembloid at day 14 of co-culture. The paraffin section was stained using hematoxylin and eosin (H&E stain). **D:** Immunofluorescence analysis of neuro-mesodermal assembloid at day 14 in co-culture. The histological section incubated with specific antibodies targeted against the endothelial marker-protein PECAM (CD31) and the neuron-specific marker-protein β-III tubulin (TUJ1). A CD31^+^ vascular network is visible at the neuro-mesodermal interface. **E:** Semiquantitative RT-PCR analyses detecting neural crest marker genes (TFAP2, Sox10), peripheral neuron marker genes (Peripherin, Phox2b, Isl1) as well as sensory neuron (Brn3a) and sympathetic neuron (TH) markers. Detection of GAPDH was used as loading control. Analyses were performed using mRNA from assembloids at different time points in coculture (day 5 – day 40).

Neural and mesodermal cell aggregates were differentiated separately from iPS cells utilizing different combinations of small molecules and cytokines as previously described ^7 8^. Neural aggregate formation takes seven days and is induced by Purmorphamin (PMA), SB431542, Dorsomorphin and CHIR99021 (Fig.1 A). Formation of neural rosettes and expression of the neuronal marker TUJ1 are hallmarks of successful differentiation (Fig.1 CD). The generation of mesodermal aggregates takes four days and is directed by CHIR99021 and BMP4. The mesodermal part is characterized by loose connective tissue and the presence of CD31-positive blood vessel-like structures which form a perineural vascular plexus at the neuro-mesodermal interface (Fig.1 C-D).

Neuro-mesodermal assembloids were kept in co-culture for up to 40 days and were initially characterized by immunofluorescence staining and RT-PCR analyses.

Semiquantitative RT-PCR revealed the expression of neural crest cell markers (TFAP2 and Sox10) as well as sensory and sympathetic peripheral neuron markers (Peripherin, Isl1, Brn3a, TH). These initial experiments gave a first hint indicating the formation of a peripheral nervous system (Fig.1E) ^9^.

Rigorous histological analyses using immunofluorescence microscopy revealed the appearance of Sox10^+^ NCC-like cells within the neural part of the assembloid at day 10 of co-culture (Fig.2 A). A part of these cells is TFAP2-positive while others appear TFAP2/Sox10 double-positive, probably representing migrating neural crest cells (Fig.2 A) ^10^. After 20 days in co-culture, Sox10^+^ neural crest cells were found to migrate towards and into the mesodermal part of the assembloid (Fig.2 B-C). Moreover, ganglion-like clusters of peripheral nerve cell perikarya were found at the interface between neural and mesodermal part (Fig.2 C-D‘). Cells within these ganglion-like structures stained double-positive for Peripherin and Isl1 (Fig.2 D’), confirming their peripheral neuron identity ^11 12^. Additionally, they were also found to be Brn3a-positive (Fig.2 E-E‘) identifying them as sensory neurons ^13^. 3D-reconstruction of confocal fluorescence microscopic images show a typical pseudounipolar cell morphology (Fig.2 F-F‘). From that, we conclude that migrating neural crest cells give rise to sensory ganglia like the dorsal root ganglion (DRG) which forms early during PNS development near the neural tube.

**Figure 2:**
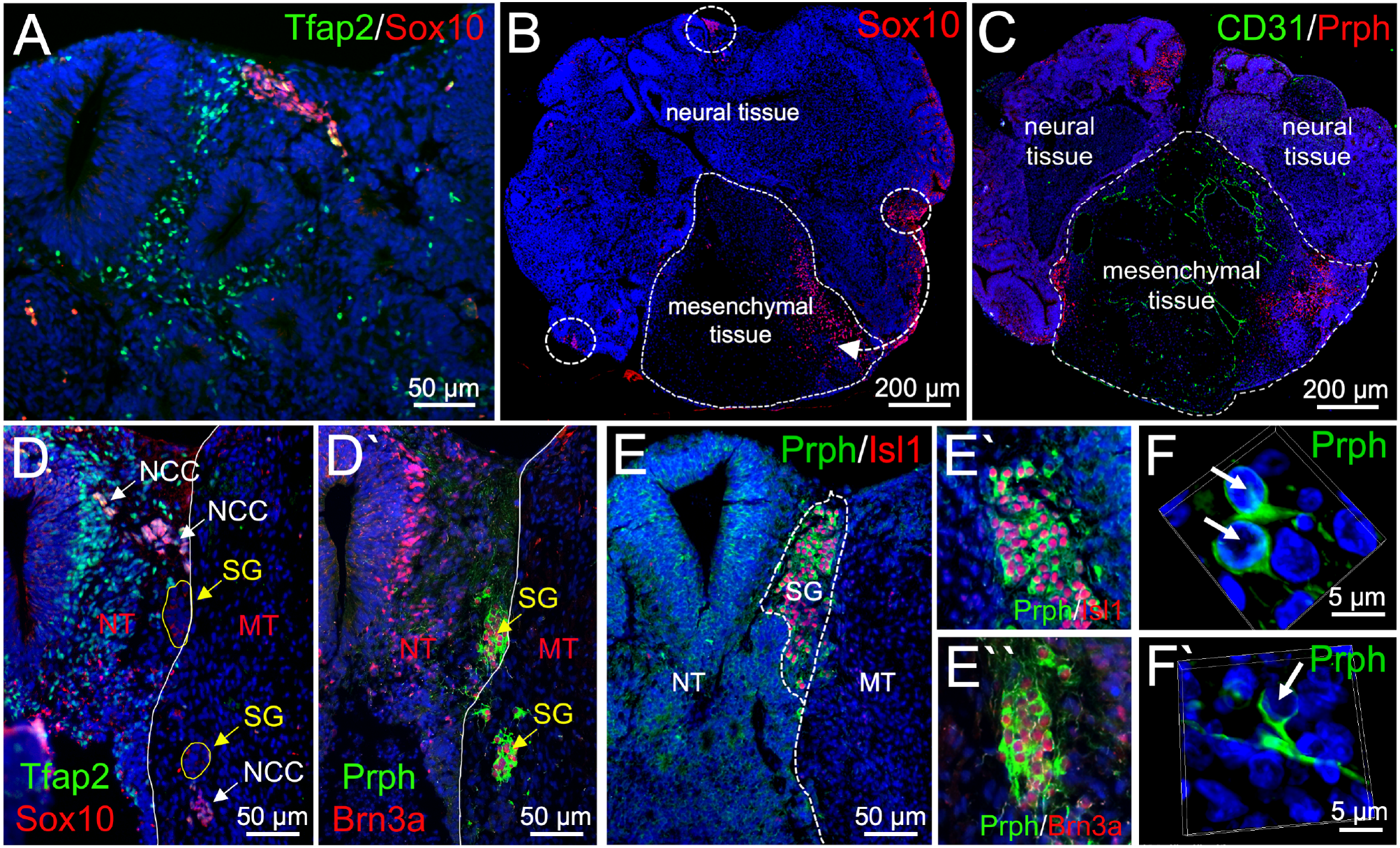
Neural crest cell migration and sensory ganglion formation in neuro-mesodermal assembloids. **A-F‘:** Immunofluorescence analysis performed on sections from neuro-mesodermal assembloids at day 14 in co-culture. **A:** Detection of TFAP2 and Sox10 expression. **B:** Detection of Sox10 expression shows neural crest migration from the neural into the mesodermal part of the assembloid. **C:** Peripherin+ cells indicating PNS-neurons are detected at the neuro-mesodermal interface. A CD31^+^ vascular plexus is visible in the mesodermal part of the assembloid. **D-D‘:** While TFAP2^+^/Sox10^+^ neural crest cells (NCC) migrate from the neural (NT) into the mesodermal (MT) part of the assembloid, Peripherin^+^/Brn3a^+^ sensory ganglia (SG) form along the migration tract near the neuro-mesodermal interface. **E-E‘:** Cells within the Sensory ganglia coexpress the sensory neuron marker proteins Peripherin, Brn3a and Isl1. **F-F‘:** Sensory neurons show typical pseudo-unipolar morphology.

Bundles of peripheral nerve fibers were found to project from the sensory ganglia into the mesenchymal tissue and were accompanied by Sox10^+^ cells (Fig.3 A-B). Moreover, 3D-reconstruction of whole mount stained, and tissue cleared assembloids revealed that peripherin-positive neuronal processes do not only grow towards the mesenchymal but also the neural part (Fig.3 C, Supplemental Movie 1) creating an interface between peripheral and central nervous system. Interestingly, peripheral nerve fibers in the mesenchymal part were accompanied by Sox10-positive cells (Fig.3 D), which was not the case in the neural part (Fig.3 D‘). This is equal to the *in vivo* situation in which peripheral neurons are myelinated by Sox10^+^ Schwann cells while central fibers are myelinated by Sox10^-^ Oligodendrocytes.

**Figure 3:**
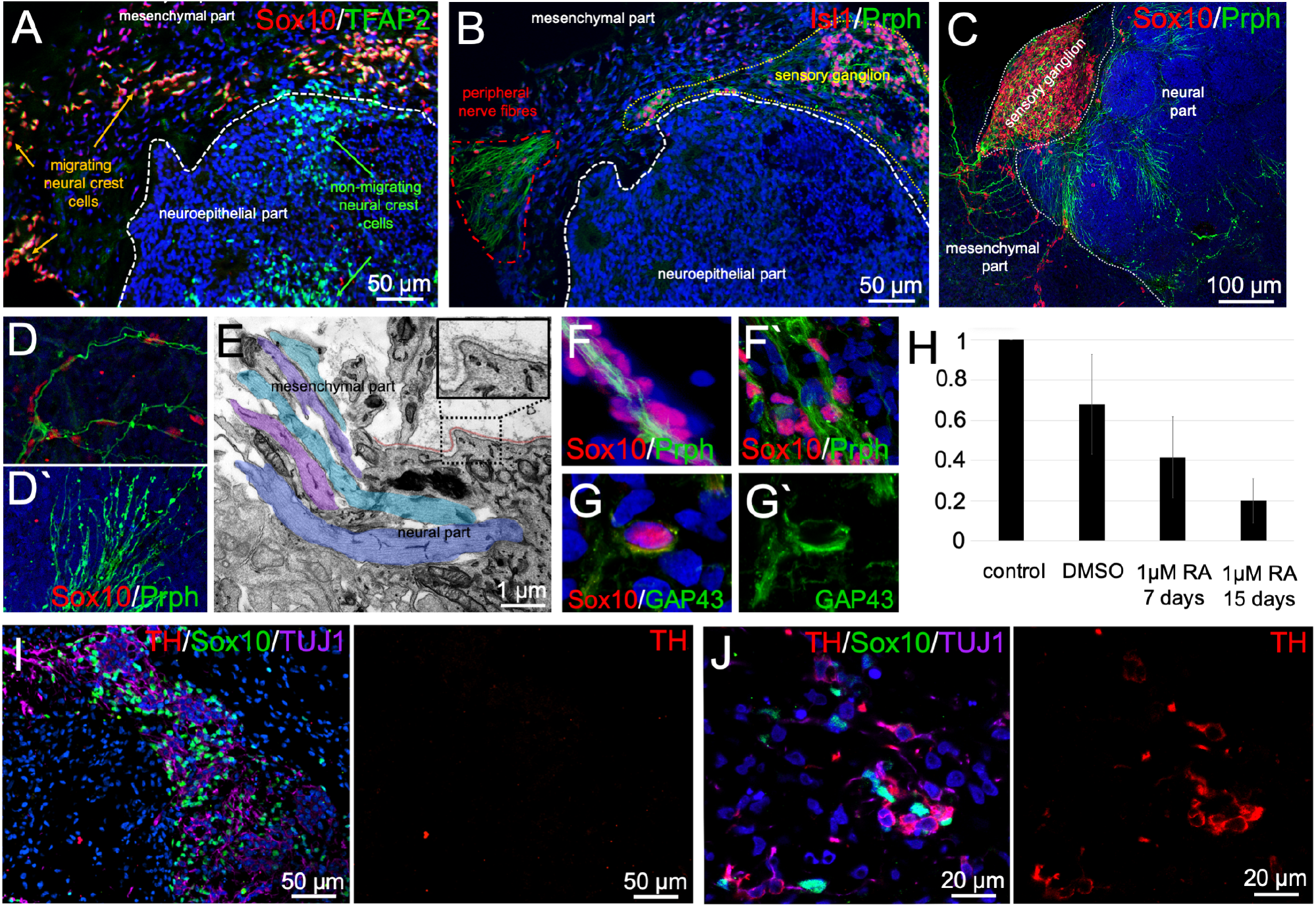
Peripheral nerve formation and appearance of sympathetic ganglia in neuro-mesodermal assembloids. **A-B:** Immunofluorescence analyses performed on 20 days old assembloids show neural crest cell migration indicated by Sox10 and TFAP2 expression (A) as well as Isl1/Peripherin^+^ ganglion formation (B). Sensory ganglia send peripheral nerve-like axon bundles into the mesodermal part of the assembloid. **C:** 3D reconstruction of immunofluorescence pictures taken from tissue cleared assembloid. A Sox10/Peripherin^+^ ganglion is depicted sending axonal projections to the neural and the mesodermal part of the assembloid. **D:** Axons projecting to the mesodermal part are accompanied by Sox10^+^ Schwann cell-like cells (D). Axons projecting to the neural tissue are devoid of Sox10^+^ cells (D’). **E:** Transmission electron microscopic (TEM) analysis reveals axons (blue, purple and green color) penetrating the basement membrane (brown color) at the neuro-mesodermal interface marking the basal side of the neuroepithelium. **F-G:** Peripheral nerve-like axon bundles are accompanied by Sox10/GAP43^+^ Schwann cell-like cells. **H:** Treatment of assembloids with 1 μM retinoic acid (RA) reduces Sox10 expression in a time dependent manner. The solvent DMSO also showed a mild effect on Sox10 expression. **I:** Large sensory ganglia at the neuro-mesodermal interface do not show expression of the sympathetic neuron marker TH. **J:** Small TH/Sox10^+^ sympathetic ganglia can be found deeper within the mesodermal part of the assembloid.

Transmission electron microscopy (TEM) revealed that bundles of axons perforate the basement membrane at the basal side of the neuroepithelium to connect neural and mesenchymal part of the assembloid (Fig.3 E). We conclude, that Sox10-positive cells accompanying peripheral nerve fiber bundles (Fig.3 F-F‘) represent Schwann cells or Schwann cell precursors, which are exclusively found in the PNS. This assumption is supported by the detected co-expression of Sox10 and GAP43 (Fig.3 G-G‘) ^14^. Interestingly, younger assembloids show bundles of axons in contact with each other but isolated from the surrounding tissues by a sheath of Schwann cells as observed in the early embryonic peripheral nerve (Fig.3 F) ^15^. Later, the Schwann cells invade the nerve, separating the axons into smaller bundles (Fig.3 F‘).

To proof that the assembloid model is suitable for drug testing applications or toxicological screenings, we treated the cultures with different concentrations of retinoic acid (RA). RA has been shown to impact neural crest cell delamination and migration and can cause a condition termed fetal retinoid syndrome which results in neural crest migration defects ^16^. We observed that RA treatment reduced the mRNA expression level of Sox10 in the assembloid cultures in a time dependent manner. The solvent DMSO showed a mild effect on Sox10 expression as well (Fig.3 H).

In the embryo, besides forming sensory ganglia, the migrating NCCs give rise to sympathetic ganglia as well. These are characterized by co-expression of Tyrosine Hydroxylase (TH) and Peripherin ^12 17^. We could not detect any TH-positive cells within the sensory ganglia (Fig.3 I) but found TH/TUJ1 double-positive cells in larger distance from the neural part forming small ganglion-like structures along with Sox10^+^ Schwann cells (Fig.3 J). This is in line with the *in vivo* situation where sensory ganglia are located near the CNS while sympathetic ganglia are located more distant within the sympathetic trunk or associated with the abdominal aorta or its major branches ^18^. This indicates, that the assembloid model recapitulates neural crest induction, delamination, migration as well as sensory and sympathetic ganglion formation.

The forming ganglia were observed to be surrounded by a dense CD31^+^ vascular network (Fig.4 A). Within the mesodermal part of the assembloid, peripheral nerve fibers get in contact with CD31^+^ blood vessels (Fig.4 B). 3D reconstruction of whole mount stained, and tissue cleared assembloids reveal a close interaction of Peripherin^+^ sensory nerve fibers with the developing vascular plexus (Fig.4 C, Supplemental Movie 2). Moreover, when the axons contact the vessels, they form bouton en passant-like structures indicating synapse formation (Fig.4 D-E, white arrow) which is not observed in axons not contacting a vessel (Fig.4 D-E, yellow arrow).

**Figure 4:**
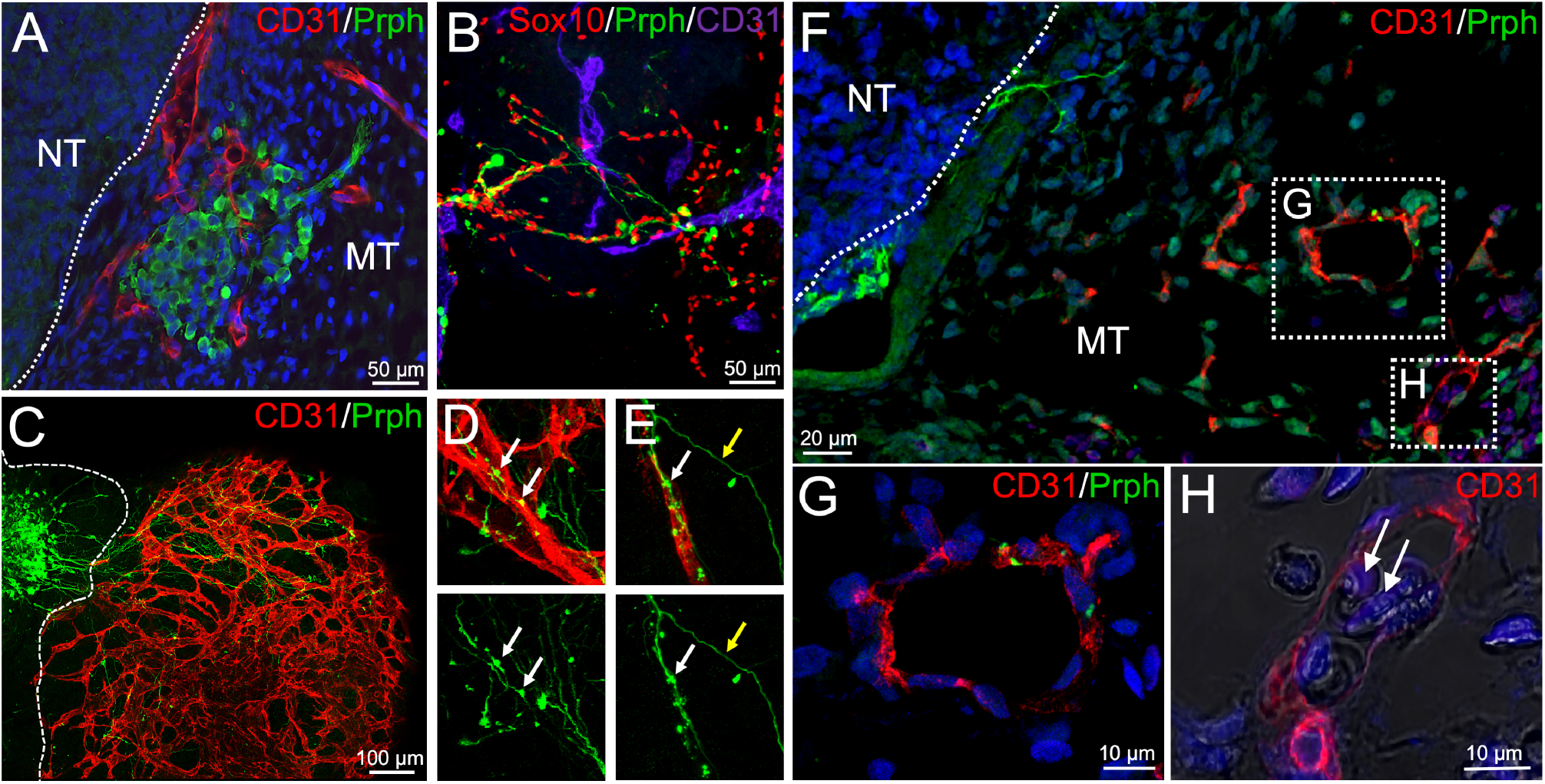
Close interaction of blood vessels and peripheral nerves in neuro-mesodermal assembloids. **A:** Immunofluorescence analyses performed on sections from neuro-mesodermal assembloids at day 20 in co-culture. Peripherin^+^ ganglia at the neuro-mesodermal interface are enwrapped by an endothelial network. **B:** Migrating neural crest cells (Sox10^+^) and peripheral neurons (Peripherin^+^) were observed to be in close contact with CD31^+^ vascular structures. **C:** 3D reconstruction of immunofluorescence pictures taken from tissue cleared assembloid. A large sensory ganglion is depicted sending multiple axonal projections into the mesodermal part of the assembloid. Moreover, a complex and well-organized vascular plexus can be observed. **D-E:** Peripheral nerve fibers align with blood vessels and form bouton en passant-like structures (white arrow). These structures are not observed in axons without vessel contact (white arrow). **F-H:** Transplantation of neuro-mesodermal assembloids on the chicken chorioallantoic membrane (CAM). 6 days after transplantation, assembloids were analyzed for Peripherin and CD31 expression (F). Peripheral ganglia and nerve fibers could be observed. Peripherin^+^ fibers were found in close contact to CD31^+^ human blood vessels (G). Chicken erythrocytes (white arrow in H) were observed inside the vessel lumen indicating vessel functionality (H).

To test the functionality of the vascular network, neuro-mesodermal assembloids were transplanted on the chicken chorioallantoic membrane, which led to the connection of human assembloid vessels to the chicken circulatory system (Fig. 4 F-H). 6 days after transplantation, paraffin sections were made, and immunofluorescence analyses were performed. We were able to show CD31^+^ vascular structures as well as Peripherin^+^ ganglia and nerve fibers. Again, peripheral nerve fibers were observed to contact human blood vessels (Fig.4 G). Moreover, nucleated chicken erythrocytes were found inside the vessel lumen indicating perfusability (Fig.4 H).

To test the functionality of sensory ganglia, we performed calcium imaging. For that purpose, assembloids were loaded with the calcium indicator Fluo4-AM and treated with the TRPV1-receptor agonist capsaicin ^19^. We observed that intracellular calcium levels increased 2-fold in cells located within sensory ganglia in response to capsaicin treatment (Fig. 5, Supplemental Movie 3).

**Figure 5:**
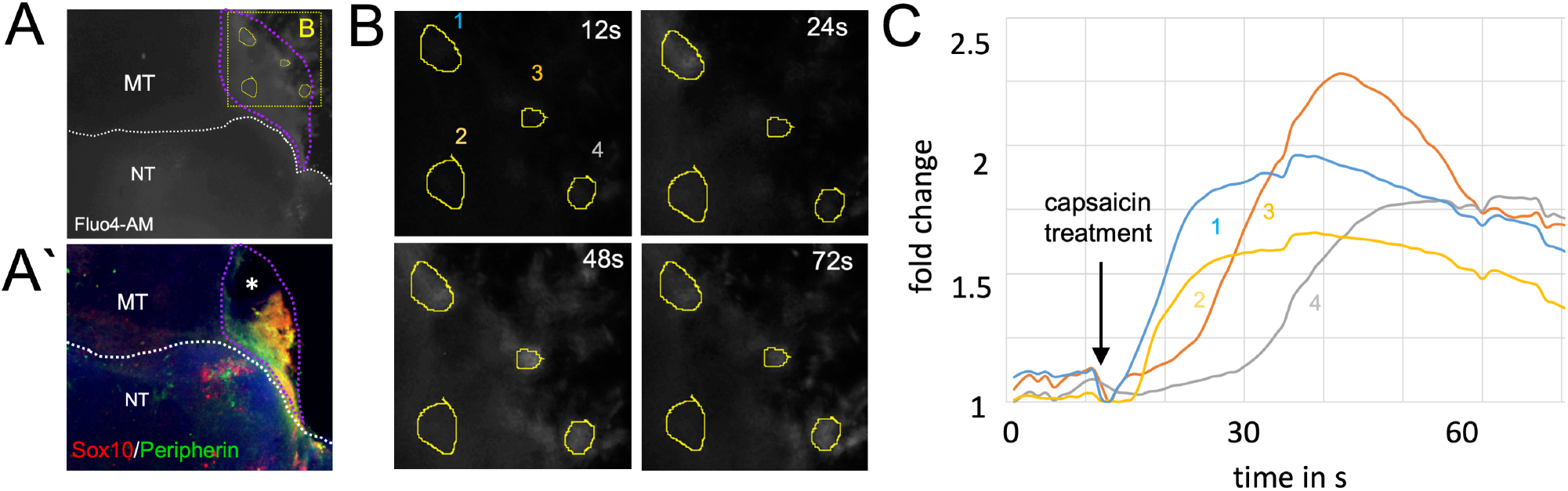
Peripheral ganglia are responsive to capsaicin treatment. **A:** Neuro-mesodermal assembloids were loaded with the calcium indicator dye Fluo4-AM and treated with capsaicin. Elevated calcium levels in response to capsaicin treatment were observed in a distinct region of the organoid (purple dotted line). See also Supplemental Movie 3. **A‘** After calcium measurement, assembloids were fixed in 4% PFA solution, whole mount stained for Sox10 and Peripherin, and tissue cleared. 3D reconstruction of confocal immunofluorescence images confirmed the location of a peripheral ganglion in the capsaicin responsive region. Tissue was partially destroyed during sample preparation at black area marked with white asterisk. **B:** Higher magnification of the image depicted in A (yellow dotted square). Pictures show Fluo4-AM fluorescence at different time points (12s, 24s, 48s, 72s). Capsaicin was added to the cultures after 12 s. **C:** Quantification of Fluo4-AM fluorescence intensity in 4 selected regions of interest (ROI) (yellow circles in image A and B)

## Discussion

Here we present a novel human *in vitro* assembloid model mimicking several aspects of peripheral nervous system development. The assembloids allow to investigate mechanisms of neural crest cell migration in 3D-cell culture without any need for animal experimentation.

The assembloids are easy to culture and develop within 20-40 days in agarose coated non-adhesive 96-well plates. They can be manufactured in large numbers with comparable differentiation outcome (Fig. S1), which makes them an attractive platform for toxicity testing or drug screenings e.g., to identify potential teratogenic substances which influence neural crest delamination and migration during embryonic development ^20^ as demonstrated in a proof-of-concept experiment using retinoic acid ^16^.

Although a variety of tissues and early developmental embryo stages have been mimicked in the lab within the last decade, 3D tissue culture models of neural crest migration and PNS development are still rare. A recent publication describes neural crest migration and PNS development in the complex environment of long-term cultured elongating human gastruloids ^21^. Two other publications describe neuromuscular organoids. These develop from neuromesodermal progenitor cells and show the formation of motorneurons and sensory neurons along with skeletal muscle ^22 23^. An interesting paper by Xiao and colleagues describes a method to directly reprogram mouse and human fibroblasts into sensory neurons. The authors observed the appearance of ganglion-like cell clusters during the reprogramming process, which they termed “sensory ganglion organoids” ^24^. However, to our opinion, these structures do not fully meet the definition of the term “organoid”.

Here we present an alternative and easy to handle neuro-mesodermal assembloid model, which is generated by co-culturing of mesodermal and neural spheroids. An interesting aspect of our model is that migrating NCCs co-develop and closely interact with a complex vascular plexus as found in the embryo ^25^. Moreover, the developing sensory ganglia are highly vascularized as also observed *in vivo* ^26^. Finally, peripheral nerve fibers are getting in close contact with the developing blood vessels and form synaptic boutons en passant. This allows a close interaction of migrating neural crest cells, sensory neuroblasts and neurons with endothelial as well as peri-endothelial cells of the vascular plexus. Such neuro-vascular interactions have been discussed to be involved in developmental processes such as arterial specification and vessel branching ^27^ or axonal guidance ^28^. Moreover, an interaction between endothelial cells and sensory neuroblasts regulates both their quiescence and differentiation behavior e.g., *via* endothelial-neuroblast cytoneme contacts and Dll4-Notch signaling ^29^. Finally, Sox10^+^ neural crest cells have been demonstrated to contribute to vascular development in multiple organs ^30^. For that reason, the presented assembloid model is a promising platform to study such neuro-vascular interactions in more detail directly in the human tissue context.

## Methods

### Cell culture and neuro-mesodermal assembloid formation

To generate neuro-mesodermal assembloids, human foreskin derived iPSCs (STEMCCA NHDF iPSCs and Sendai NHDF iPSCs) were used as previously described ^7^. iPS cells are cultured on human ESC Matrigel (Corning) coated 6 well plates in StemMACS iPS Brew XF (human) medium (Miltenyi Biotec) supplemented with 10 mM ROCK inhibitor Y-27632 (Miltenyi Biotec) until they reach a confluency of 80 %. Subsequently, cells are dissociated by using StemProAccutase cell detachment solution (Gibco/Life Technologies) and gentle pipetting.

For the induction of both, neural and mesodermal organoids, cells are resuspended in StemMACS iPS Brew medium supplemented with 10 mM ROCK inhibitor Y-27632 and 4 × 10^3^ cells (in 100 μl medium) are seeded per well of a 96 well plate. The 96-wells were previously coated with 1 % Agarose (Biozym) to keep cells in suspension. Cells are cultured for 24 h at 37°C and 5% CO_2_ in a humidified incubator to allow aggregation.

To induce neural differentiation, iPSC aggregates are cultured in neural induction medium 1 (NIM1) (Neurobasalmedium (Gibco) 50 %, DMEM-F12 (Gibco) 50 %, B27 without Vitamin A (Gibco) 1 x, N2-Supplement (Gibco) 1 x, L-Glutamine (Gibco) 2 mM, CHIR 99210 (Sigma-Aldrich) 3 μM, SB431542 (Miltenyi Biotec) 10 μM, Dorsomorphin (Tocris) 1 μM, Purmorphamine (Miltenyi Biotec) 0.5 μM) for 48 h. Subsequently the medium is changed to neural induction medium 2 (NIM2) (Neurobasalmedium 50 %, DMEM-F12 50 %, B27 without Vitamin A 1 x, N2-Supplement 1 x, L-Glutamine 2 mM, CHIR 99210 3 μM, Ascorbic acid (Sigma-Aldrich) 0.06 mg/ml (355 μM), Purmorphamine 0.5 μM) for 72 h. After the induction phase, medium is changed to neural differentiation medium (NDM) (Neurobasalmedium 50 %, DMEM-F12 50 %, B27 without Vitamin A 1 x, N2-Supplement 1 x, L-Glutamine 2 mM, Ascorbic acid 0.06 mg/ml (355 μM), Penicillin/Streptomycin (Sigma-Aldich) 1 x) for 24 h.

For mesodermal induction, iPSC aggregates are cultured in mesoderm differentiation medium (MDM) (Advanced DMEM-F12 (Gibco) 100 %, L-Glutamine 2 mM, Ascorbic acid 0.06 mg/ml (355 μM), CHIR 99210 10 μM, BMP4 (Pepro Tech) 25 ng/ml) for 72 h.

Finally, a single neural and a single mesodermal aggregate per agarose coated 96-well are brought in co-culture to allow assembloid formation. Neuro-mesodermal assembloids are kept in suspension culture in NDM for up to 40 days for 24 h at 37°C and 5% CO_2_ in a humidified incubator.

### Immunofluorescence analyses

Co-cultured organoids are fixed in 4 % paraformaldehyde (PFA) solution (Sigma-Aldrich) overnight and subsequently washed with PBS (Sigma-Aldrich). Afterward, 5 μm paraffin (Merck) sections were prepared. For immunofluorescence analyses, sections were deparaffinized, rehydrated and unmasked using Sodium Citrate buffer (10 mM, pH6). Primary antibodies to TUJ1 (Biozol, 801202), CD31 (DAKO, M0823), SOX10 (R&D Systems, AF2864), Peripherin (Merck-Millipore, AB1530), TFAP2 (DSHB, 3B5), Isl1 (DSHB), GAP43 (Novos Biology, NB300-143), TH (Abcam, AB112) and Brn3a (Santa Cruz, SC-8429) were diluted in NBS blocking solution and incubated overnight at 4 °C. Secondary Cy2-, Cy3- or Cy5-labelled antibodies (Dianova) diluted in PBS were used to visualize primary antibodies. Secondary antibodies were incubated for 1 h at room temperature. To highlight the tissue structure, paraffin sections were further H&E-stained utilizing Eosin y-solution (AppliChem) and Gill‘s Hematoxylin solution (Chroma). Images were taken using an Axiovert 40 CFL microscope (Zeiss) and an Eclipse Ti confocal laser scanning microscope (Nikon).

### Whole mount staining and tissue clearing

For tissue clearing the organoids were fixed in 4 % PFA in PBS overnight at 4 °C. Afterwards, organoids were washed 3 × 30 min in PBS. Subsequently, organoids were incubated in an ascending MeOH (Sigma-Aldrich) series (30 min 50 % MeOH in PBS, 30 min 80 % MeOH in PBS, 2 × 30 min 100 % MeOH) and washed 2 × 30 min in 20 % DMSO (Carl Roth) in MeOH. Next, a descending MeOH series followed (30 min 80% MeOH in PBS, 30 min 50 % MeOH in PBS, 30 min PBS). Subsequently, specimens were incubated 2 × 30 min in 0.2 % Triton X-100 (Sigma-Aldrich) in PBS and kept in penetration buffer (PBS, 0.2 % Triton X-100, 0.3 M Glycine (Carl Roth), 20 % DMSO) at 37 °C overnight. After that, organoids were incubated in blocking solution (PBS, 0.2 % Triton X-100, 0.3 M Glycine, 6 % BSA, 10 % DMSO) overnight at 37 °C. For the staining process, organoids were washed 2 x for 1 h in washing buffer (0.2 % Tween-20 (ApplyChem) in PBS) and afterwards incubated with primary antibodies in blocking buffer (PBS, 20 % Tween-20, 4 % BSA, 5 % DMSO) for 24 h at 37 °C. Following primary antibodies were used: CD31 (DAKO, M0823), Sox10 (R&D Systems, AF2864), and Peripherin (Merck-Millipore, AB1530). The organoids were then washed twice for 30 min in washing buffer and afterwards incubated with secondary antibodies overnight at 37 °C in blocking solution. Finally, organoids were washed 3 × 30 min in washing buffer and dehydrated in an ascending MeOH series (3 × 30 min in 50 % MeOH in PBS, 3 × 30 min in 80 % MeOH in PBS and 3 × 30 min in 100 % MeOH). For tissue clearing, samples were incubated in ethyl cinnamate (Sigma-Aldrich) for 24 h at 37 °C.

### Transmission electron microscopy

Organoids were fixed in 2.5 % glutaraldehyde, 4 % PFA in 0.1 M cacodylate buffer (50 mM cacodylate, 50 mM KCl, 2.5 mM MgCl2, pH7.2) on ice for 2 h and washed 4 x with 0.1 M cacodylate buffer, 5 min each. Organoids were subsequently fixed for 60 min with 1 % osmium tetroxide in 0.1 M cacodylate buffer and washed 2 x in 0.1 M cacodylate buffer, 10 min each. After washing with bi-distilled water, organoids were dehydrated in an ascending EtOH series using solutions of 30 %, 50 %, 70 % EtOH, 10 min each. Organoids were contrasted with 2 % uranyl acetate in 70 % EtOH for 60 min and subsequently dehydrated using an EtOH array of 70 %, 80 %, 90 %, 96 % and two times 100 % for 10 min each. Subsequently, organoids were incubated in propylene oxide (PO) two times for 30 min before incubation in a mixture of PO and Epon812 (1:1) overnight. The following day, samples were incubated in pure Epon for 2 h and embedded by polymerizing Epon at 60 °C for 48 h.

Specimen were cut using an ultramicrotome, collected on nickel grids, and post-stained with 2.5 % uranyl acetate and 0.2 % lead citrate. Finally, specimens were analyzed using a LEO AB 912 transmission electron microscope (Zeiss).

### Quantitative Real Time PCR

For quantitative real-time PCR, RNA of 30 days old organoids is extracted by using the Direct-Zol™ RNA MiniPrep Plus kit (Zymo Research) following manufacturer‘s instructions. Complementary DNA was prepared by using GoScript Reverse Transcriptase™ (Promega). Quantitative PCR analyses are conducted using the GoTaq® Q-PCR Master Mix (Promega) and a StepOnePlus™ Real-Time QPCR thermocycler (Applied Biosystems).

The following primer pairs were used:

GAPDH: FW 5’-TGACAACTTTGGTATCGTGGA-3’ and RV 5’-CCAGTAGAGGCAGGGATGAT-3’,
SOX10: FW 5’-CCTCACAGATCGCCTACACC-3‘ and RV 5‘-CATATAGGAGAAGGCCGAGTAGA-3’.

### Semiquantitative RT-PCR Analyses

For semiquantitative RT-PCR analyses, RNA is extracted by using the Direct-Zol RNA MiniPrep Plus Kit (Zymo Research). To generate cDNA, the GoScript Reverse Transcriptase (Promega) is used as described in the manufacturer‘s protocol. The polymerase chain reaction was performed using the Red MasterMix (2x) Taq PCR MasterMix (Genaxxon) using the following primer pairs:

Brn3a: FW 5’-GGGCAAGAGCCATCCTTTCAA-3’ and RV 5’-CTGTTCATCGTGTGGTACGTG-3’,
GAPDH: FW 5’-TGACAACTTTGGTATCGTGGA-3’ and RV 5’-CCAGTAGAGGCAGGGATGAT-3’,
Isl1: FW 5’-GCGGAGTGTAATCAGTATTTGGA-3’ and RV 5’-GCATTTGATCCCGTACAACCT-3’,
Peripherin: FW 5’-GCCTGGAACTAGAGCGCAAG-3’ and RV 5’-CCTCGCACGTTAGACTCTGG-3’,
PHOX2b: 5’-AACCCGATAAGGACCACTTTTG-3’ and RV 5’-AGAGTTTGTAAGGAACTGCGG-3’,
SOX10: FW 5’-CCTCACAGATCGCCTACACC-3’ and RV 5’-CATATAGGAGAAGGCCGAGTAGA-3’,
TFAP2: FW 5’-AGGTCAATCTCCCTACACGAG-3’ and RV 5’-GGAGTAAGGATCTTGCGACTGG-3’,
TH: FW 5’-GGAAGGCCGTGCTAAACCT-3’ and RV 5’-GGATTTTGGCTTCAAACGTCTC-3’

### Calcium Imaging

Calcium imaging was performed on organoids glued to a cover slip using 20 μl human ESC Matrigel. Organoids were loaded with 1 μM Fluo4-AM (Gibco) and incubated at 37 °C for 30 min. Subsequently, organoids were washed with PBS and kept in culture medium. To test the functionality of sensory ganglia, capsaicin (Thermo Fisher Scientific), dissolved in Chloroform, was added to culture medium to a final concentration of 1 μM. A video of the dynamic fluorescence changes is recorded using a Leica DM IL LED microscope equipped with an external light source for fluorescence excitation (Leica EL6000) and a Leica DFC450C camera. Quantification and data analysis was performed using Image J (Fiji) and Microsoft Excel.

### Chick chorioallantois membrane (CAM) assay

Fertilized chicken eggs were incubated at 37 °C at a humidity of 60 %. After 5 days of incubation, assembloids are transplanted on the CAM of the developing chicken embryo. For transplantation assembloids are attached to a round nylon mesh (diameter of appx. 5 mm; 150 μm grid-size, 50% open surface, 62 μm string diameter, 35 g/m^2^, PAS2, Hartenstein) with 10 μl Matrigel. After gelling at 37°C, the mesh is placed upside down on the CAM. After 5-10 days, the mesh with the organoids is carefully explanted, washed in PBS and fixed for 1 h in 4 % PFA. Subsequently, 5 μm paraffin sections are prepared and processed for immunofluorescence analyses as described above. All procedures comply with the regulations covering animal experimentation within the EU (European Communities Council DIRECTIVE 2010/63/EU). They are conducted in accordance with the animal care and use guidelines of the University of Würzburg.

## Acknowledgements

We thank Doris Dettelbacher-Weber, Martina Gebhardt, Erna Kleinschroth, Karin Reinfurt-Gehm, Ursula Roth, Sieglinde Schenk, Elke Varin and Lisa Wittstatt for excellent technical assistance and all members of the Stem Cell and Regenerative Medicine Group for their support. This work was supported by grants of the IZKF-Würzburg (Interdisziplinäres Zentrum für Klinische Forschung der Universität Würzburg) (project E-D-410) to P.W. and the German Research Foundation (DFG) by the Collaborative Research Center TR225-B04 to S.E..

## Author Contributions

P.W. conceived the study and designed experiments; A.F.R., and P.W. conducted experiments; N.W.: performed electron microscopic analyses; S.E.: acquired funding and provided feedback and expertise; A.F.R. and P.W. wrote the manuscript and performed final approval. All authors have read and agreed to the published version of the manuscript.

## Supplemental Figures

**Figure S1:**
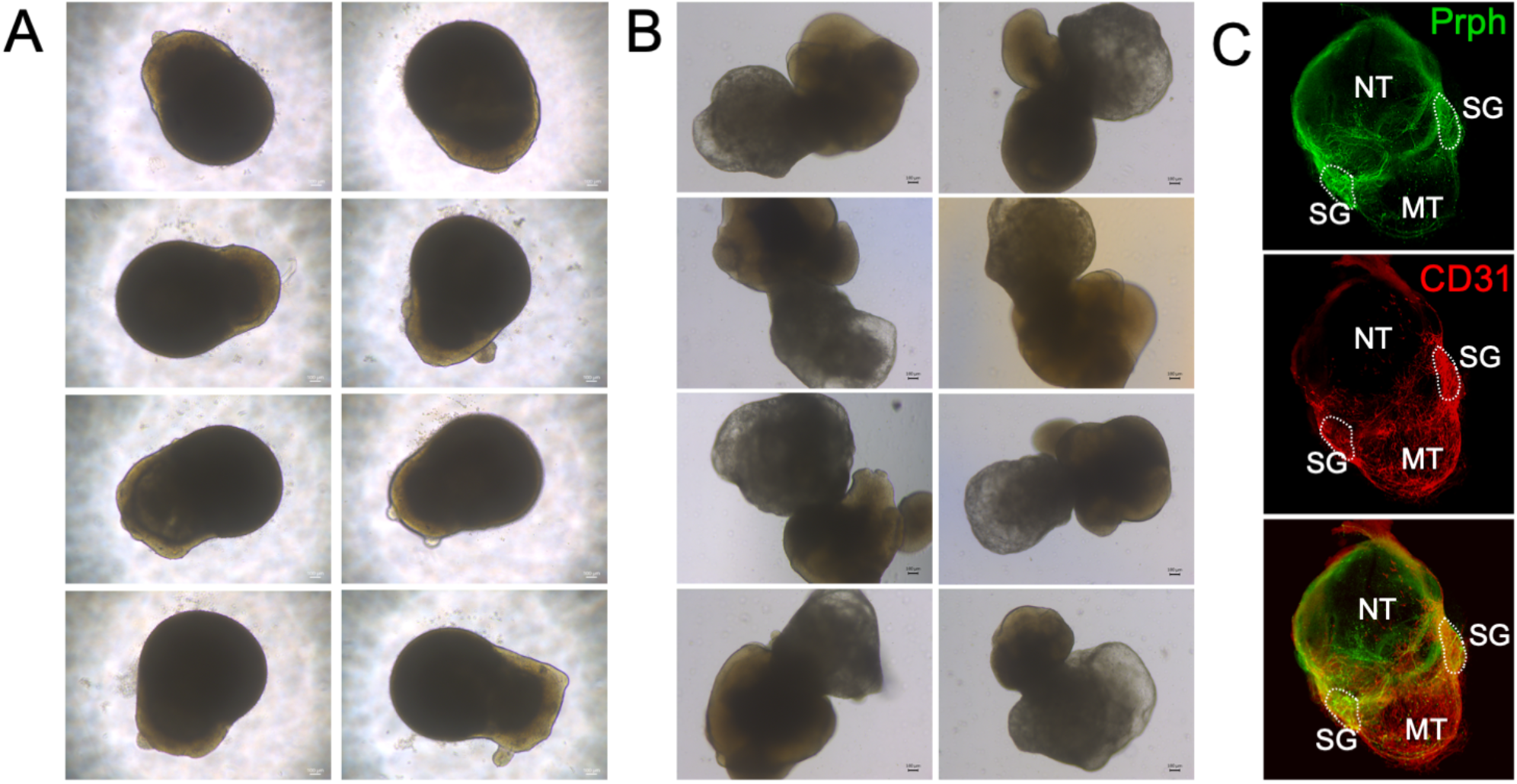
Reproducibility of the experimental procedure. **A:** 8 individual representative organoids at day 7 in co-culture are depicted. **B:** 8 representative neuro-mesodermal organoids at day 14 in co-culture are depicted. **C:** Whole mount immunofluorescence analyses using specific antibodies targeted against CD31 and Peripherin (Prph) were performed. Images show a 3D reconstruction of confocal microscopic images from a representative tissue cleared organoid (SG: sensory ganglia, NT: neural tissue, MT: mesodermal tissue).

## Supplemental Movies

**Movie S1: Large sensory ganglion at the neuro-mesodermal interface**

The movie shows a large sensory ganglion at the neuro-mesodermal interface. Peripheral neurons (green) were detected using specific Peripherin antibody. Neural crest cells and derivates were detected using a Sox10 antibody (red). Nuclei are stained with DAPI. The movie shows a 3D reconstruction from a z-stack of confocal images taken by laser scanning microscopy.

**Movie S2: Blood vessels and peripheral nerves in the mesodermal part of the assembloid**

The movie shows a hierarchically organized network of blood vessels (red) and peripheral nerve fiber bundles (green) in the mesodermal part of the assembloid. Peripheral neurons were detected using specific Peripherin antibody. Blood vessels were detected using a CD31 antibody. The movie shows a 3D reconstruction from a z-stack of confocal images taken by laser scanning microscopy.

**Movie S3: Sensory ganglion cells show increased calcium levels in response to capsaicin treatment**

The movie shows a calcium imaging experiment (left side). The intracellular calcium levels were visualized using Fluo4-AM. Calcium levels rise in response to capsaicin treatment. Video plays at 8x speed. After calcium imaging, the assembloid was fixed and whole mount stained using specific antibodies targeted against Peripherin (green) and Sox10 (red) to visualize the position of the peripheral ganglion (right side).

